# scRNA-Seq-based drug repurposing targeting idiopathic pulmonary fibrosis (IPF)

**DOI:** 10.1101/2022.09.17.508360

**Authors:** Anika Liu, Joo-Hyeon Lee, Namshik Han, Andreas Bender

## Abstract

Idiopathic pulmonary fibrosis (IPF) is a chronic lung disease, which affects around three million people worldwide and is characterized by impaired regeneration from recurrent injury to the alveolar epithelium resulting in progressive lung scarring. In this work, we target a cell transition in the differentiation of AT2 cells into mature AT1 cells which is inhibited in IPF and contributes to the regeneration of the alveolar epithelium. We hypothesize that inducing this transition promotes lung regeneration and can ameliorate the disease. To this end, we characterized the intermediate population to AT1 cell transition signature using multiple recently generated single-cell RNA-Seq datasets on murine bleomycin injury and IPF patients. We then matched this signature to drug perturbation signatures retrieved from the LINCS database to identify the most suitable candidates for drug repurposing. Compound classes identified in this analysis include glucocorticoids, HSP90 inhibitors, HDAC inhibitors and a range of kinase inhibitors. Furthermore, we are able to interpret our findings by identifying multiple potential direct targets and downstream effectors, such as NFKB1 and HIF1A. Overall, we show how scRNA-Seq data can potentially be leveraged for drug repurposing by enabling better characterization of cell transitions which can subsequently be used for signature matching.

## Introduction

Idiopathic pulmonary fibrosis (IPF) is a form of chronic interstitial lung disease (ILD) which affects ∼3 million people worldwide and shows increasing incidence rates, also because it primarily affects elderly adults. It is characterized by progressive lung scarring, thickening of the interstitium and continuous decline in lung function, which results in shortness of breath, also known as dyspnoea, and cough. This severely affects the patient’s quality of life and a poor prognosis with a median survival rate of 3-5 years if untreated ^1^.

Currently, two treatments are approved for IPF: Nintedanib ^2^, which inhibits tyrosine kinases involved in proangiogenic and pro-fibrotic pathways, and pirfenidone ^3^, which has anti-fibrotic, anti-inflammatory, and antioxidant activity. Both small molecules effectively reduce disease progression and decline of pulmonary function but are not able to improve lung function or quality of life ^4^. While severe side effects are rare, drug discontinuation due to adverse events was still found in 20.9% of subjects on pirfenidone and 26.3% on nintedanib ^5,6^. This leaves lung transplantation as the only cure and only alternative intervention, but this is only possible for a minority of patients due to the limited organ availability as well as comorbidities and age of the patients ^7^. Overall, this combination of a large and growing patient population, severe and deadly disease burden, and limited availability of treatment options makes IPF a societal unmet medical need ^8,9^.

Promoting endogenous alveolar repair has emerged as a promising therapeutic strategy, which may not only slow down or stop disease progression but can potentially even cure IPF by regenerating lung function ^10^. Recently, an intermediate progenitor cell state was identified in AT2 to AT1 differentiation which expresses epithelial and mesenchymal markers, as well as markers of senescence such as TGFβ which indicates a pro-fibrotic role ^11–13^. This transitional cell state was also found to persist in fibrotic regions of IPF lungs, and explicitly around foci of high collagen expression ^11,12^. This suggests that the terminal differentiation from this transitional cell state to functional AT1 cells is inhibited and that it contributes to fibrogenesis.

In the murine bleomycin lung injury model, which is one of the most extensively used and best-characterized preclinical models for acute and chronic lung injury ^14^, the AT2 to AT1 trajectory was further characterized using scRNA-Seq identifying an intermediate stem cell state expressing similar markers, including pro-fibrogenic factors and distinct cell-cell communication with mesenchyme and macrophages ^15,16^. Additionally, chronic inflammation by sustained IL-1b treatment was found to inhibit the terminal differentiation to functional AT1 cells, also resulting in an accumulation of intermediate progenitors ^16^.

Promoting the intermediate progenitor to AT1 transition is hence therapeutically interesting from two angles: To restore the depleted AT1 population which may help to restore lung function, and to reduce the level of pro-fibrotic intermediate progenitors.

Single-cell transcriptomics has fundamentally revolutionised biological research by enabling us to study gene expression at cellular resolution instead of averages across samples. With this technology, it is now possible to study questions on cellular heterogeneity and compositional changes, and to identify rare cell populations which may be overlooked in bulk transcriptomics ^17,18^. Furthermore, transitional cell states and cell trajectories can be uncovered with scRNA-Seq, e.g. in perturbation response or differentiation, overall providing us with a better understanding on how dynamic cellular processes are orchestrated ^17^.

Single-cell transcriptomics is already advancing our systems-level understanding of diseases ^19,20^ and treatments ^21,22^, which will likely impact the development of future therapeutics. However, we believe that scRNA-Seq cannot only guide drug development indirectly through mechanistic insights, but that the data itself can also be exploited to prioritize compounds in drug discovery. The approach we revisit in this study towards this aim is signature matching to identify compounds inducing perturbation responses which match a defined disease signature. Previous studies applying signature matching to scRNA-Seq data follow the idea of reversing the diseased state to individual cell clusters instead of bulk expression, which mitigates compositional changes as covariate ^23,24^. Matching results are then available for each cluster, and a final drug ranking can then be obtained, e.g. by giving higher priority to compounds detected in multiple clusters, which was implemented by Wang *et al*. ^23^, or by summarising to an overall score, as implemented in ASGARD (A Single-cell Guided pipeline to Aid Repurposing of Drugs) ^24^.

While the assumption made in these approaches that compounds targeting more cell clusters are more promising is plausible in the absence of further information, we often know which cell types are causally involved in pathogenesis and can use this to direct the analysis. For example, Alakwaa ^25^ focussed on type II alveolar cells (AT2 cells) in their repurposing study on COVID-19, or more precisely they used the differential expression comparing ACE2-expressing AT2 cells and other AT2 cells as disease signature because ACE2 was already hypothesized to be a receptor for severe acute respiratory syndrome coronavirus 2 (SARS-CoV-2) at the time of the study ^97^. While this hence includes prior information on relevant cell types, the underlying hypothesis that reducing the expression of the target receptor may help to treat COVID-19 is not further elaborated and not an established strategy to tackle infectious diseases.

Instead, inducing disease-relevant transitions poses a more promising application of signature matching, although not purely data-driven, given that it is then known that 1) the transition can take place which is not always true, e.g. due to lineage commitment ^26^, and that 2) inducing the transition is affecting pathogenesis and e.g. not merely reversing symptoms. This has already been successfully applied in previous bulk transcriptomics studies, e.g. osteoblast differentiation has been targeted to identify treatments for osteoporosis ^27,28^, and candidates for differentiation therapy have been proposed based on the transition signature of leukaemia cells to granulocytes ^29^.

With bulk transcriptomics, however, it was only possible to characterise the transitions for particular cell types *in vitro*, which limits the physiological relevance, while transitions were difficult to characterize *in vivo* given expression changes cannot be assigned to particular cell population and rare cell types may be overlooked completely. However, with the advent of scRNA-Seq, it is now possible to discover and characterize disease-relevant and rare cell types and transitions *in vivo* which opens up new opportunities, which to our knowledge have not been explored yet.

In this study, we focus on the transition from a rare disease-enriched intermediate progenitor cell state, recently discovered aided by scRNA-Seq ^12,13^, to AT1 cells, which are the alveolar epithelial cells involved in gas exchange in the lung. While studies in the bleomycin injury model have already elucidated some signals involved in the AT2 to AT1 differentiation further in mice ^11,15,16^, specific proteins or pathways with established ability to induce the transition have yet to be uncovered in IPF suspending target-based drug discovery. Using a signature matching-based approach as alternative, we are able to leverage perturbation signatures from LINCS to prioritize compounds for drug repurposing as regenerative medicine. As summarised in Figure 1 and further described in Methods, we first derive transition signatures and consensus compound perturbation signatures for this which are then matched. Additionally including compound bioactivity data from ChEMBL^30^, this is further interpreted through the identification of target proteins, and a target miRNA, which may be causally linked to the cell transition.

**Figure 1:**
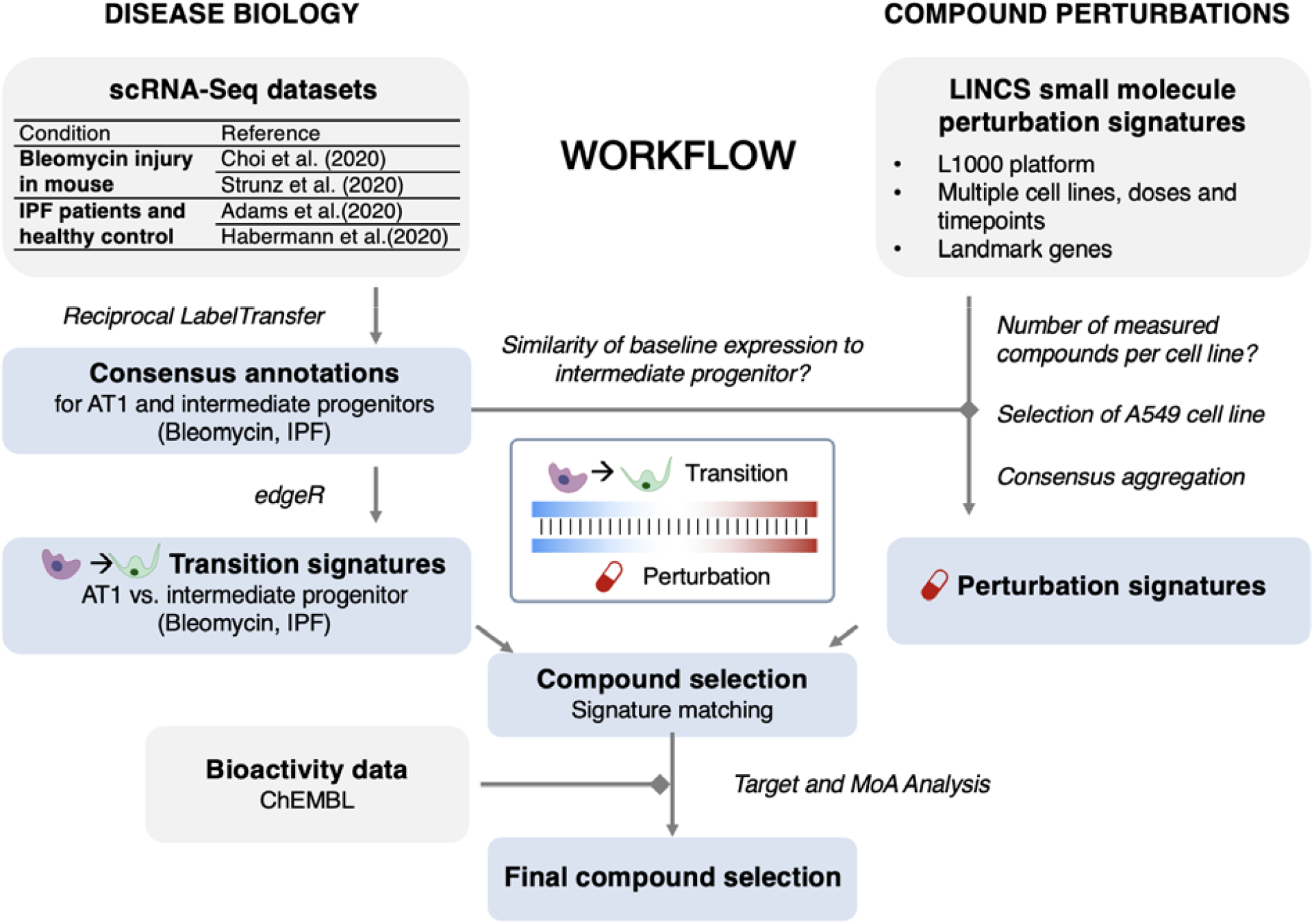
General workflow of the analysis. We first characterised the transition signatures in IPF, as well as in bleomycin-induced pulmonary fibrosis in mice which is a commonly used animal model for IPF. Based on these, we match perturbations which induce similar transcriptional changes, identifying multiple distinct compound classes. To further interrogate the potential targets and mechanisms, we additionally take advantage of bioactivity data from ChEMBL^30^.

## Methods

### Deriving transition signatures from scRNA-Seq data

In this study, we used scRNA-Seq data from two studies on bleomycin injury in mice and two studies on IPF patients, respectively (Table 1). For these datasets, count matrices as well as metadata were derived via the GEO FTP server for all datasets except for the metadata by Choi *et al*. ^16^ which was provided directly by the authors instead (File S1). Throughout the study, the single-cell data was handled in `Seurat` ^31^. From the study by Habermann *et al*. ^12^, only samples from IPF patients and healthy controls were used although the study covered multiple lung diseases, in order to avoid potential confounding factors. For each sample with at least 1,000 cells, doublets were removed using `scDblFinder` ^32^ which generated artificial doublets between randomly selected cells based on the 1,000 top expressed genes and subsequently classified cells as doublets if a large number of artificial doublets were identified in its neighbourhood taking into account an expected doublet rate of 1% per thousand cells captured with an uncertainty of 40%. The classifier was retrained three times removing cells labelled as doublets after each iteration. Subsequently, all samples from the same publication were merged, normalized and variance stabilized using `SCTransform` ^33^. PF datasets were subset to epithelial cells based on the original annotations as these can be clearly distinguished and are at the centre of this study.

**Table 1:** scRNA-Seq dataset origins.

| Paper | Condition | Cells | GEO ID | Platform |
| --- | --- | --- | --- | --- |
| **Choi *et al.***<sup>16</sup> | Bleomycin injury in mouse | Alveolar epithelium | GSM4304609, GSM4304611, GSM4304613 | 10x Genomics Chromium |
| **Strunz *et al.***<sup>15</sup> | Bleomycin injury in mouse | Epithelium | GSE141259 | Dropseq |
| **Adams *et al.***<sup>13</sup> | IPF patients and healthy control | Whole lung | GSE136831 | 10x Genomics Chromium |
| **Habermann *et al.***<sup>12</sup> | PF patients and healthy control | Whole lung | GSE135893 | 10x Genomics Chromium |

### Harmonize cell annotations through reciprocal label transfer

To identify a consensus cell annotation for both PF and both bleomycin datasets, respectively, we then predicted for each dataset, referred to as “query”, whether each cell is labelled as an AT1 cell, intermediate progenitor or another cell type based on the cell annotations from the respective other dataset, referred to as “reference” as described by Stuart *et al*. and implemented in `Seurat` ^34^. First, we first identified pairs of cells between two datasets which are mutual nearest neighbours (MNN) using the *FindTransferAnchors* function. These transfer anchors were among each other’s 5-nearest neighbours based on the PCA reduction of the reference (30 PCs) and its projection onto the query dataset, and were further scored and filtered using default parameters. Subsequently, the *TransferData* function was used to construct a weight matrix which quantifies the association between each query cell and each anchor, which was then multiplied with the anchor classifications to compute the label predictions. We then only included AT1 cells and intermediate progenitors in the downstream analysis which were regarded as such based on both the original and the predicted annotation.

Details on annotation concordance are visualized in Figure S 1 using an integrated UMAP representation for PF and bleomycin, respectively, derived through `Harmony` ^35^. For the bleomycin datasets, the integration (and hence the visualization) included cell types which are expected to be present in both datasets, namely AT1, AT2 and intermediate progenitor cells, to account for the fact that Choi *et al*. specifically studied cells originating from AT2 cell lineage. In contrast, all epithelial populations were included in PF except for ciliated cells which formed a distinct cluster. Overall, good agreement was found between the cross-predicted cell labels and the original annotations of each dataset with the same label (AT1, DATP or other) being achieved in 84.1% and 97.4% for the datasets by Strunz *et al*. and Choi *et al*., respectively, while even better agreement was found for the IPF datasets by Adams *et al*. (98.5%) and Habermann *et al*. (99.1%). The confusion matrices in Figure S 1C additionally show that there are intermediates progenitors which were rather regarded as AT1 by Strunz *et al*. than by Choi *et al*. and vice versa. As such annotation discrepancies can affect how well a joint signature can be derived, only consensus intermediate progenitors and AT1 cells, which were regarded as such based on the original and predicted annotation, were included in the downstream analysis.

### Differential expression and disease signature analysis

Based on the derived consensus annotations, transition signatures were derived by comparing AT1 cells to intermediate progenitors in IPF or bleomycin injury, respectively, using `edgeR` ^36^ implementation of the likelihood ratio test (LRT) as part of the `Libra` package ^37^. In this pseudo-bulk approach, we specified individual samples as replicates which translates to the individual timepoints in case of bleomycin injury or individual patients in case of IPF. For downstream analysis, the derived bleomycin signature was mapped from mouse gene symbols to HGNC symbols using `biomaRt` ^38^. For Reactome pathway maps ^39^ and miRNA target genesets ^40^, derived using `msigdbr` ^41^, enrichment was computed using the `fgsea` package ^42^ for all gene sets with 10-200 genes per geneset in order to capture processes which can be statistically inferred and are biologically informative. Signalling pathway activity was inferred with `PROGENy` based on the expression of the top 100 genes involved in the pathway at hand according to significance ^43^. To gain further insights into proteins involved in upstream signalling, TF activity was inferred with the *msviper* function from the `viper` package ^44^ for all `DoRothEA` ^45^ regulons with confidence levels A-C for which at least 10 genes were measured.

Subsequently, we inferred upstream signalling networks using `CARNIVAL 46` based on the inferred TF activities as measurements, a signed and directed protein-protein interaction network from Omnipath^47^ as prior knowledge, and CPLEX ^48^ as ILP solver. Default parameters for this were loaded using the *defaultCplexCarnivalOptions* function.

### Signature matching

Baseline expression profiles for the cancer cell lines used in LINCS was obtained from the Cancer Cell Line Encyclopedia ^49^ (CCLE) through the depmap portal (21Q3, DOI: `10.6084/m9.figshare.15160110.v2`). For the intermediate progenitor populations, the baseline expression in TPM (transcripts per million) was determined in `Seurat` based on the consensus labelled cells ^31^.

LINCS drug perturbation signatures (Level 5) from the alveolar A549 cell line were obtained from GEO using GEO ID GSE92742. All signatures from the same compound, dose and time combination were combined into a single consensus signature using moderated z-scoring which was first introduced for summarising biological replicates from the same experiment in the LINCS analysis pipeline ^50^ and was also implemented to summarise multiple signatures from the same condition across experiments by Szalai *et al*. ^51^. As the 978 landmark genes are chosen based on complementarity and other genes are inferred based on these, only the measured the genes are used for signature matching while all genes are used to infer TF activity using `DoRothEA` ^45^ using the same settings as for the disease signatures.

Pearson correlation between each drug and disease signature was computed using `cTRAP` ^52^, which provides multiple measures used in signature matching. While significant (p-value < 0.05) and positive Pearson correlation were required for both transition signatures, the magnitude of Pearson correlation was subsequently used to further reduce the list of drug repurposing candidates for experimental validation.

### Target bioactivity

To derive bioactivity data on the perturbations from CHEMBL 30 ^30^, we first used the Python client for the ChEMBL API ^53^ to identify molecules with the same connectivity based on the canonical SMILES representation, provided as metadata in LINCS, and subsequently retrieved all pCHEMBL values for these compounds where the target organism was *Homo sapiens*. Besides the target metadata from CHEMBL, additional information on the protein families of the targets was obtained from the Uniprot knowledgebase (UniProtKB) ^54^, and the UniProt ID was also used to map targets to gene symbols using BioMart ^38^ via the `biomaRt` R package.

To compare bioactivity across compounds, we used pCHEMBL values, which are defined as -Log(molar IC_50_, XC_50_, EC_50_, AC_50_, K_i_, K_d_ or Potency), and hence enables integration of multiple metrics of half-maximal response ^30^. To establish enrichment of target activity for matched compounds, we define pCHEMBL values > 5 as “active” while pCHEMBL values > 5 are regarded as “inactive”. We then used a one-sided Fisher’s Exact test, as implemented in the `stats` package, and compare to unmatched compounds with perturbation signatures in the A549 cell line. Furthermore, we tested for significant correlation between pCHEMBL and the strength of signature matching, estimated by the Pearson correlation, using the `correlation` R package ^55^. In both, the correlation and enrichment analysis, the maximal signature matching correlation per disease signature was used to summarise signature matching across LINCS profiles on the same compound, and only targets with at least 20 data points were included. Baseline expression for targets was derived, and genes with a minimal TPM of 0.5 were regarded as expressed as the same cut-off was used in the Expression Atlas ^56^.

## Results and discussion

### Transcriptional characterization of the transition from intermediate progenitor to AT1

To characterize the transition signature from intermediate progenitors to AT1 cells, we first harmonized the cell annotations through reciprocal label transfer and subsequently derived differential expression signatures comparing AT1 cells to intermediate progenitors (Figure 2A and B). In these signatures, up-regulation indicates higher expression levels in AT1 compared to intermediate progenitors and, vice versa, down-regulation indicates higher expression intermediate progenitors compared to AT1 cells. We first investigated the expression of previously identified markers for AT1 cells and intermediate progenitors as quality check for the signatures and markers (Figure 2). In both signatures, we find up-regulation for all previously identified AT1 markers, such as advanced glycosylation end-product specific receptor (*AGER*), podoplanin (*PDPN*), and HOP homeobox (*HOPX*), except for insulin-like growth factor binding protein 2 (*IGFBP2*). Interestingly, *IGFBP2* was previously found to be expressed later in AT1 differentiation than other markers, such as *HOPX*, and at this earlier IGFBP2^-^ cell state AT1 cells can transdifferentiate to AT2 cells in alveolar regeneration, while IGFBP2^+^ AT1 cells do not show cellular plasticity physiologically or in injury ^57,58^. Hence, this indicates that the PF transition signature is not capturing this terminal step of AT1 differentiation, potentially because the AT1 IGFBP2^-^ cell are enriched or because mature IGFBP2^+^ AT1 cells are destroyed in IPF lungs, or because they have been favoured in the cell annotation in comparison to bleomycin injury.

**Figure 2.**
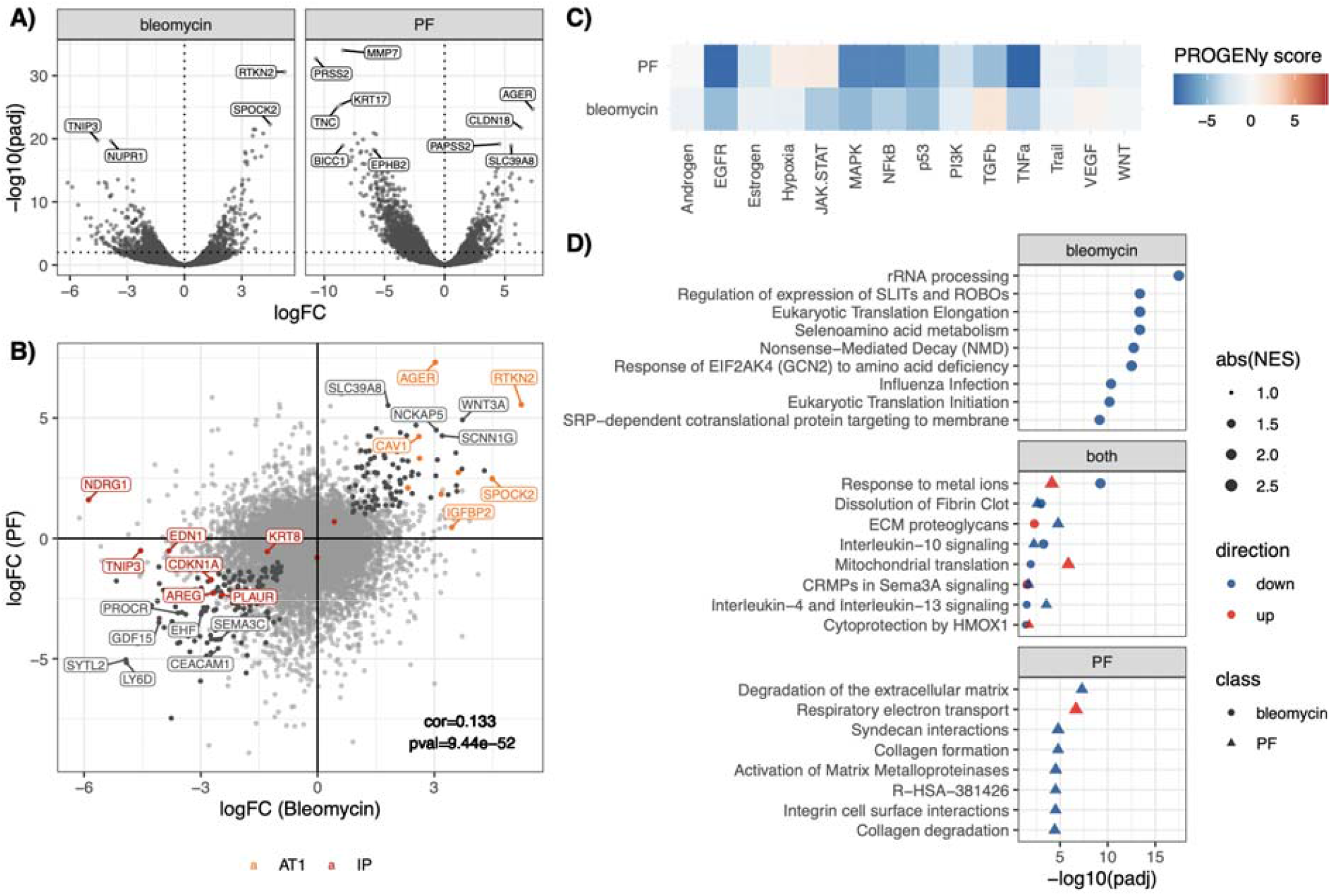
Transition signatures in bleomycin injury and pulmonary fibrosis (PF). A) Differentially expressed genes (DEGs) in bleomycin and PF signatures. B) Comparison of DEGs identifying significant but low correlation. C) PROGENy identifies consistent and strong (absolute score >3) down-regulation of EGFR, MAPK, NFκB, p53, and TNFα signalling D) Reactome pathway enrichment identifies most significant down-regulation of stress response pathways in bleomycin injury, and extracellular matrix- and mitochondria-related processes in PF. Significant down-regulation in both is found for signalling via interleukins 4,10 and 13, as well as dissolution of fibrin clots. Opposite directionality is found for multiple stress-related and developmental processes.

For the intermediate progenitor markers, decreased expression in both signatures is found for amphiregulin (*AREG*), urokinase plasminogen activator surface receptor (*PLAUR*) and cyclin dependent kinase inhibitor 1A (*CDKN1A*). For four additional markers, significantly decreased expression was only observed in bleomycin injury indicating that these are not applicable to IPF, potentially because of differences between the respective intermediate progenitor populations, e.g. linked to species differences, or their annotations (Figure S 1).

Globally comparing the transition signatures derived for bleomycin injury in mice and PF in humans, we find a significant but low correlation between the differential expression profiles indicating that the transition signatures are similar but that there are also discrepancies between both conditions which we aimed to investigate further (Figure 2A). Subsequent gene set analysis revealed that the most strongly significantly dysregulated pathways in bleomycin injury are linked to increased expression of ribosomal proteins, which are generally involved in the translation of mRNA to proteins (Figure 2D). They epigenetically regulate to protein translation through so-called “heterogeneous ribosomes” ^59^, have previously been linked to cell fate decisions ^60^, such as proliferation, differentiation or tumorigenesis, and increased expression of ribosomal proteins was found to be a characteristic of stem cells ^61^. Consequentially, higher expression in intermediate progenitors is expected and absence thereof in PF is indeed pointing towards less differentiation between the defined AT1 and intermediate progenitor populations there.

In contrast, the most pronounced pathways in PF indicate increased levels of proteins linked to the extracellular matrix (ECM), including both collagens and matrix metalloproteinases which degrade collagen, as well as interactions on the cell surface mediated by syndecans and integrins which mediate cell adhesion to the extracellular matrix via collagen ^62^ (Figure 2D). In pulmonary fibrosis, simultaneously increased collagen production and degradation are known, however, effectively resulting in collagen accumulation which is a known driver of fibrosis via TGFβ signalling ^63^. Absence of significant changes in these processes in the bleomycin injury model may be attributed to the fact that the intermediate progenitor population is less profibrotic, potentially because injury is acute and not chronic, or also due to the known differences in collagen regulation between mice and humans ^64^.

Only few pathways are found to be dysregulated in bleomycin injury and IPF (Figure 2D). These point to higher expression in intermediate progenitors of genes involved in the dissolution of fibrin clots, which is consistent with the role of fibrinolysis modulation via *PLAUR* in alveolar repair ^65^, and for inflammation-related pathways linked to interleukins IL-4, IL-10 and IL-13. Consistent with that an increased activity of NFκB and TNFα signalling is found, as well as of p53, which generally points to stress response, and of MAPK and EGFR signalling indicating proliferation (Figure 2C). However, also pathways with opposite directionality were found such as ECM proteoglycans, mitochondrial translation, HMOX1 and metallothioneins, as well as hypoxia TGFβ, VEGF and JAK-STAT signalling in PROGENy (Figure 2D). Overall, this sheds further light on the similarities and discrepancies between both conditions.

To further study which TFs are potentially involved in the transition from intermediate progenitor to AT1 cells in bleomycin injury and IPF, we inferred their activity based on regulon expression (Figure 3). Consistent down-regulation in AT1 cells compared to intermediate progenitors is observed for a strongly interlinked set of TFs, amongst others, related to stress, inflammation and proliferation with the strongest dysregulation being found for Jun proto-oncogene (JUN) and SMAD family member 3 (SMAD3). Furthermore, the strongest TF which is activated in bleomycin injury but reduced in IPF is PR-domain containing protein 14 (PRDM14) which was previously identified as an epigenetic regulator of pluripotency in mice ^66^. In contrast to that, a higher activation in IPF but lower activation in bleomycin injury is observed for MYC proto-oncogene (MYC) and zinc finger protein 263 (ZNF263), highlighting differences in transcriptional regulation between both conditions.

**Figure 3:**
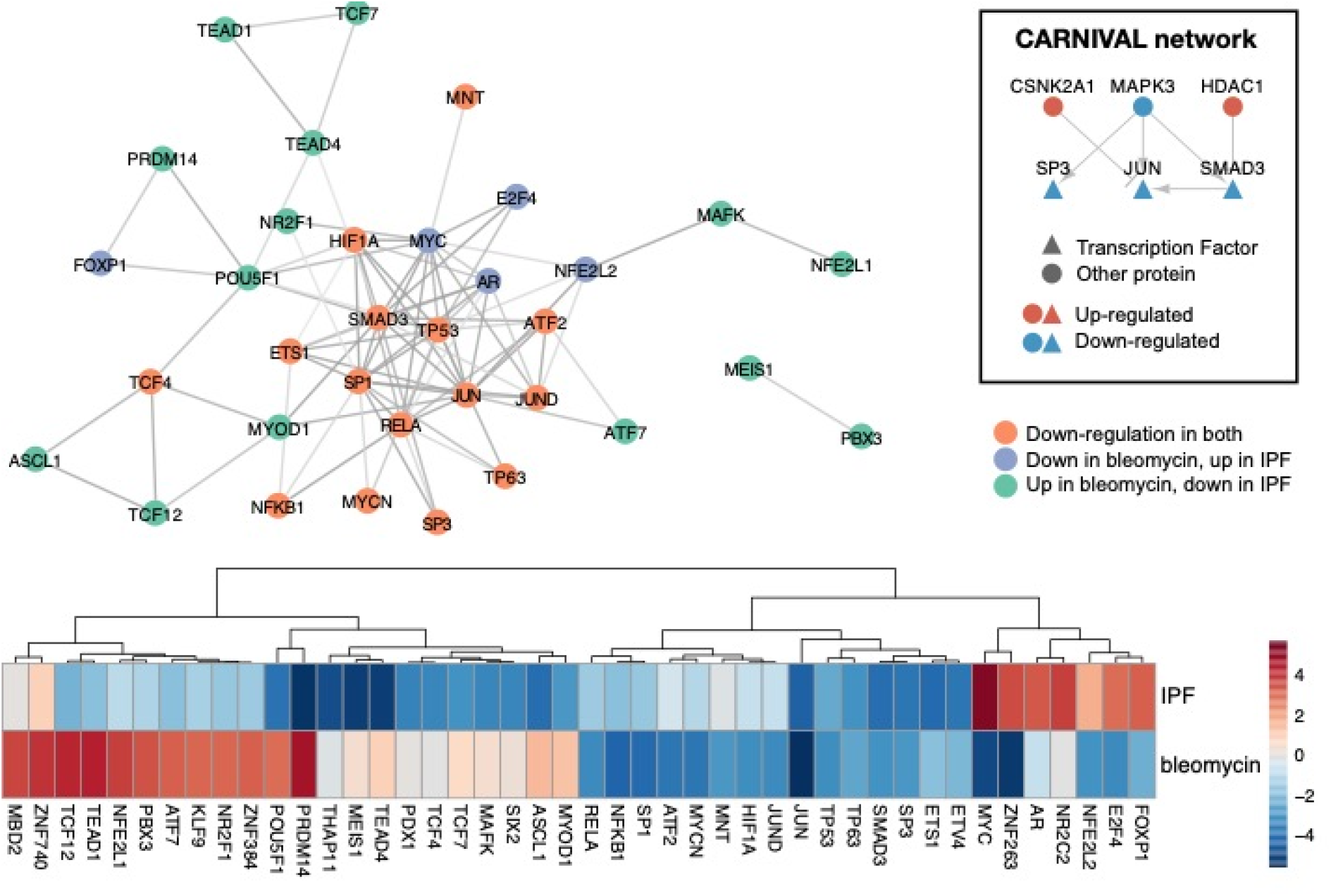
Transcription factors (TFs) involved in transition signature. TFs inferred with DoRothEA regulons confidence A-C and an absolute normalized enrichment score (NES) > 3 in at least one transition signature are shown revealing shared down-regulation for some, e.g. JUN, HIF1A and TP53, while others show opposite directionality, such as PRDM14 or MYC. Protein-protein associations with high confidence in STRING (>0.7) reveal that the core cluster largely consists of consistently down-regulated TFs, while oppositely regulated ones form separate clusters.

To further investigate potential upstream signals, we additionally inferred causal upstream networks based on both transition signatures, identifying a consistent subnetwork which consisted of the down-regulated TFs JUN, SMAD4 and SP3 as TFs as well as their potential upstream regulators. These included the down-regulated MAPK3, as well as the up-regulated histone deacetylase 1 (HDAC1) and casein kinase 2 alpha 1 (CSNK2A1) shedding further light on potential upstream signalling involved in the transition.

### Identification of repurposing candidates by signature matching

In order to identify compounds with the ability to induce transition-related gene expression changes in intermediate progenitors, we then obtained cell line perturbation signatures from the A549 cell line from LINCS due to the high number of measured perturbations and the comparably high similarity in baseline expression to the intermediate progenitor population (Figure S 2). These have been previously used as *in vitro* model for AT2 cells ^67,68^, however, do not show all characteristics of AT2 cells, such as the expression of surfactant protein or the same phospholipid content ^69,70^. We derived consensus perturbation signatures and matched these to the transition signatures for bleomycin injury in mice and IPF in humans, respectively. Overall, this identified 89 perturbation signatures mapping to 84 compounds were found to significantly correlate (p-value<0.05) with the bleomycin and PF transition signatures (Figure 4).

**Figure 4.**
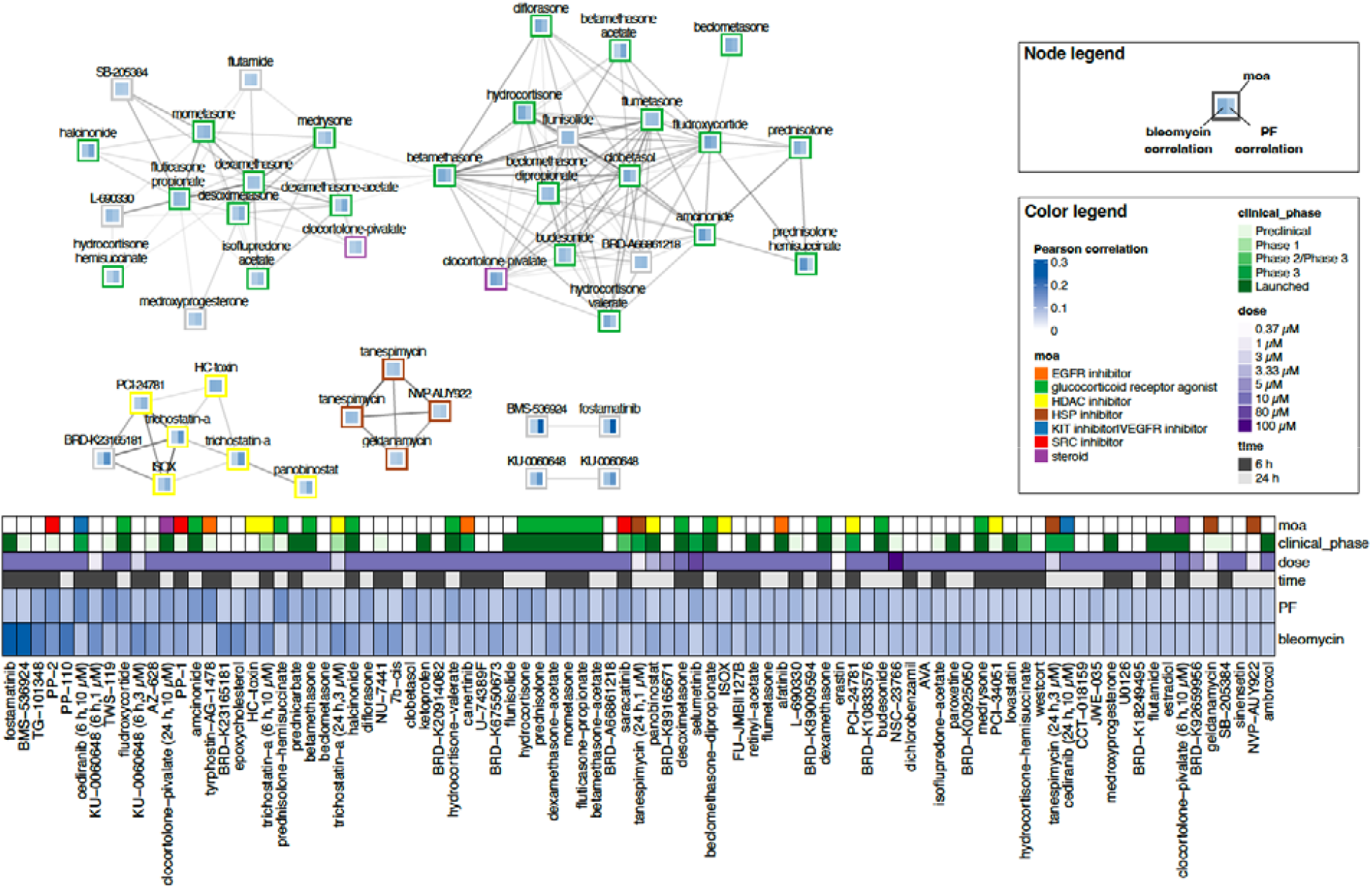
Perturbations matched to the bleomycin and PF transition signatures. The ranking by mean correlation is shown in the heatmap with the highest average correlation being observed for fostamatinib and the lowest one for ambroxol. Multiple transcriptionally similar compound clusters were identified based on the Pearson correlation between the perturbation signatures (Pearson correlation > 0.6), namely HSP90 inhibitors, HDAC inhibitors and a large cluster including glucocorticoid receptor agonists and steroids.

We next clustered the matched perturbation signatures based and identified multiple multiple compound classes, shown in Figure 4 and Figure S 3. The largest cluster contained corticosteroids, including fludroxycortide and halcinonide, which are generally anti-inflammatory and immunosuppressive ^71^, and have been widely used to treat IPF ^72^. However, despite decades of use their efficacy in IPF remains questionable as no evidence was found for an association with improved clinical outcome ^73,74^. Also multiple kinase inhibitors with overall weaker correlated perturbation signatures were found (Figure S 3). This includes the highest-ranking compound by average correlation to both signatures, the spleen tyrosine kinase (SYK) inhibitor fostamatinib which an approved treatment for chronic immune thrombocytopenia. While this has not been studied in the context of IPF, it was also prioritized in a phenotypic screen for acute lung injury ^75^ and was found to improve the clinical outcome in COVID-19 ^76^. Also nintedanib, one of the approved treatments for IPF, is in fact a tyrosine kinase inhibitor and is thought to be anti-fibrotic by targeting vascular endothelial growth factor (VEGFR), platelet-derived growth factor receptors (PDGFR) and fibroblast growth factor receptors (FGFR) ^77^. However, as kinases take in broad functions in cellular signalling, these cannot be reduced to a particular mechanism of action. Further clusters include inhibitors targeting histone deacetylases (HDACs), such as trichostatin A, and the molecular chaperone HSP90, such as tanespimycin, respectively (Figure 4 and Figure S 3). Both have not been explored heavily as targets in IPF, but have shown positive impacts on pulmonary fibrosis in animal studies. Histone deacetylases (HDACs) can acetylate of histones and other proteins and hence can epigenetically and post-translationally modulate processes ^78^. Hyperacetylation through HDAC inhibitors was found to alter gene expression in a tumour-suppressive manner countering the aberrantly high expression of HDACs generally found in cancers, and multiple HDAC inhibitors have hence been established as treatments for neoplastic diseases ^79^. Due to their antifibrotic properties, they have also been studied in the context of pulmonary fibrosis ^80^, and trichostatin A (TSA), one of the matched HDAC inhibitors, was previously found to prevent pulmonary fibrosis in rats ^81^. Furthermore, pirfenidone, one of two approved treatments for IPF with yet unknown mechanism of action, was found to modulate HDAC expression and increase histone acylation ^82^. HSP90 can support folding of damaged proteins as well as their proteasome-mediated degradation ^83^. HSP90 inhibitors were found to be efficacious in treating cancer, as HSP90 stabilizes multiple oncogenic proteins and in particular kinases, and in treating viral infections as reduced protein folding results in reduced viral activity ^84^. Also the TGFβ receptor is a client protein of HSP90 and disruption of TGFβ signalling via HSP90 inhibition was found to ameliorate bleomycin-induced pulmonary fibrosis in mice ^84,85^. Overall, this shows that compound classes are identified for which a role in pulmonary fibrosis is already being investigated or established, although not primarily due to their effects on alveolar regeneration.

### Deconvolution of potential targets and downstream TFs

We next aimed to gain further insights into the potential mechanism of action through which drugs may induce the cell transition, which may also identify new targets not yet considered for targeting alveolar regeneration and pulmonary fibrosis. We first investigated which of the TFs involved in the disease transition (Figure 3) are in fact modulated in the matched conditions and may hence be downstream effectors mediating drug action. To this end, TF activity inferred from the perturbation signatures was compared between matched signatures and signatures characterising unmatched compounds (FDR < 0.01). This identifies the most significant enrichment for the Jun Proto-Oncogene (JUN), followed by the NF-κB subunits encoded by RELA and NFKB1, all of which are down-regulated by matched compounds (Figure 5). Indeed, NKFB1 ^86^ and HIF1A ^87^ have previously been implicated in alveolar regeneration, and Choi *et al*. demonstrated that the transient activation of both is essential for the AT2 to AT1 transition ^16^. In contrast, e.g. the tumor proteins TP53 and TP63 are not differentially modulated in matched signatures, indicating that these are not targeted by matched compounds, despite similar dysregulation in the disease signature.

**Figure 5:**
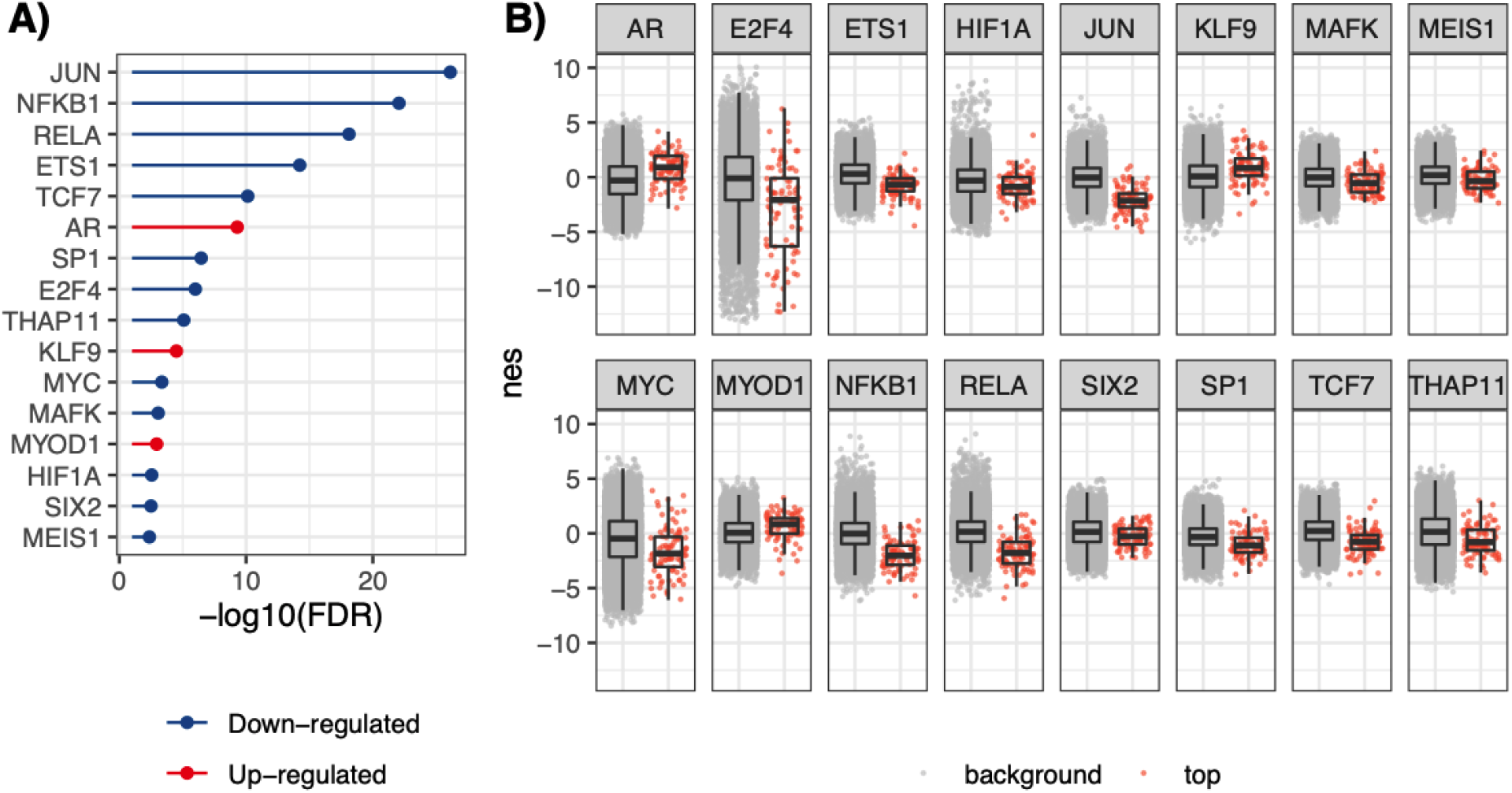
Transcription Factors (TFs) linked to disease signatures with differential regulon activity in matched LINCS signatures. Among TFs with strong dysregulation in at least one of the disease signatures (|NES| > 3), the ones with differential regulon activity between matched perturbation signatures and signatures on unmatched compounds were identified using a t-test (FDR < 0.01). The most significant differential TF activity was found for JUN, followed by RELA and NFKB1 which are downregulated in matched signatures.

To identify potential direct targets involved in the induction of the intermediate progenitor to AT1 cell differentiation, we then combined *in vitro* target activity from CHEMBL and signature matching correlation which serves as a proxy for activity on the cell transition. We first identified targets with significantly enriched activity among matched compounds (p-value) where activity is defined as pCHEMBL ≥ 5 corresponding to at least half-maximal activity at 10 μM, which was the most frequently used concentration in the matched perturbation signatures. Next, we computed the Pearson correlation between bioactivity, estimated as pCHEMBL, and maximal signature matching correlation per compound and identified six targets with significant positive correlations for both transition signatures (p-value < 0.05) which are shown in Figure 6: The glucocorticoid receptor NR3C1, which showed the strongest correlation between target activity and signature matching (Figure 7) and is known to be involved in alveolar maturation in lung development ^88^, the TFs NFKB1 and HIF1A, which were also identified in Figure 5 to show down-regulation in matched compounds based on regulon activity, and the tyrosine kinases LCK, FYN and SRC. While the matched compounds include multiple NR3C1 agonists one SRC inhibitor (PP-2), the other targets are not reported as primary mechanism of action for matched compounds. This shows that including the ChEMBL bioactivity data provides additional information on potential direct targets, and that this (unintended) polypharmacology of the compounds may contribute to their prioritization for repurposing.

**Figure 6:**
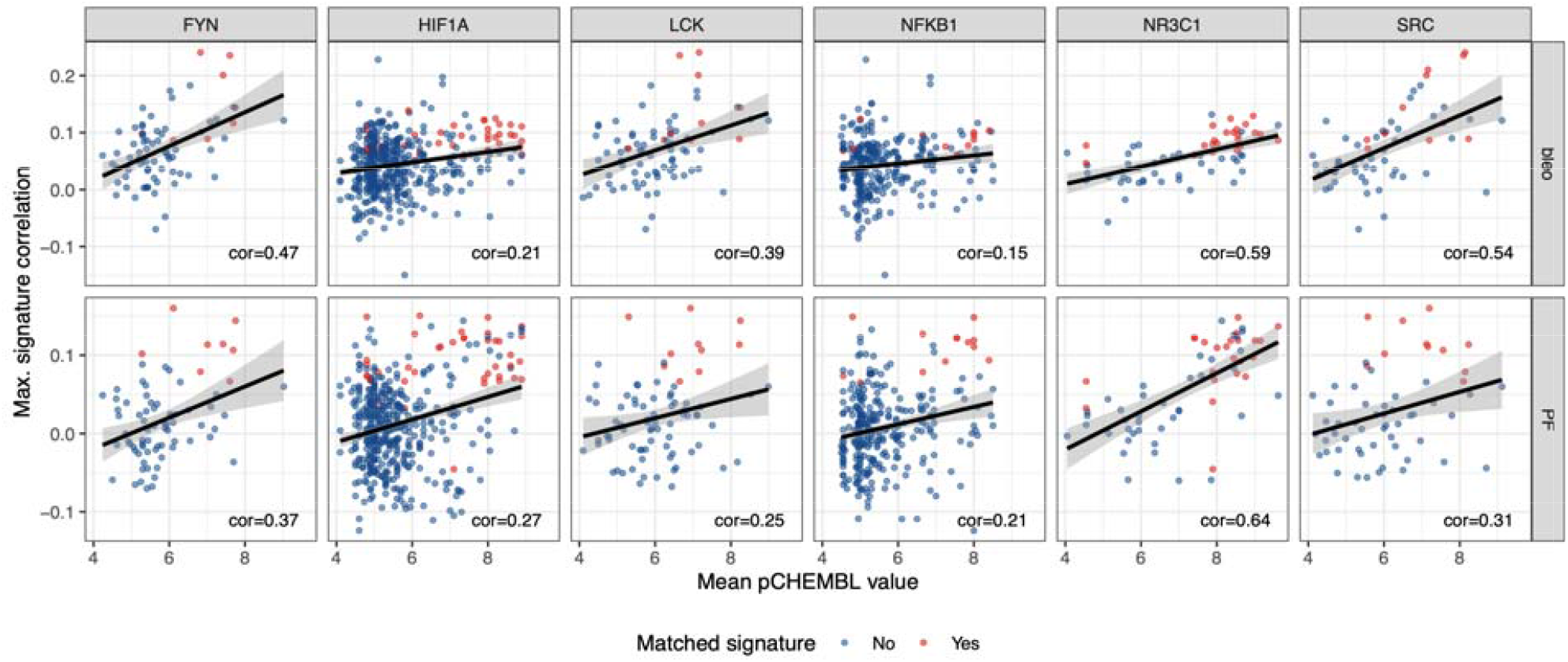
*In vitro* targets correlated with transcriptional matching. Six protein targets were identified with significant and positive correlations between pCHEMBL values and signature matching correlation for both IPF and bleomycin, namely Nuclear factor NF-κB p105 subunit (NFKB1), Hypoxia-inducible factor 1 alpha (HIF1A), glucocorticoid receptor (NR3C1), and the tyrosine kinases LCK, FYN and SRC.

**Figure 7:**
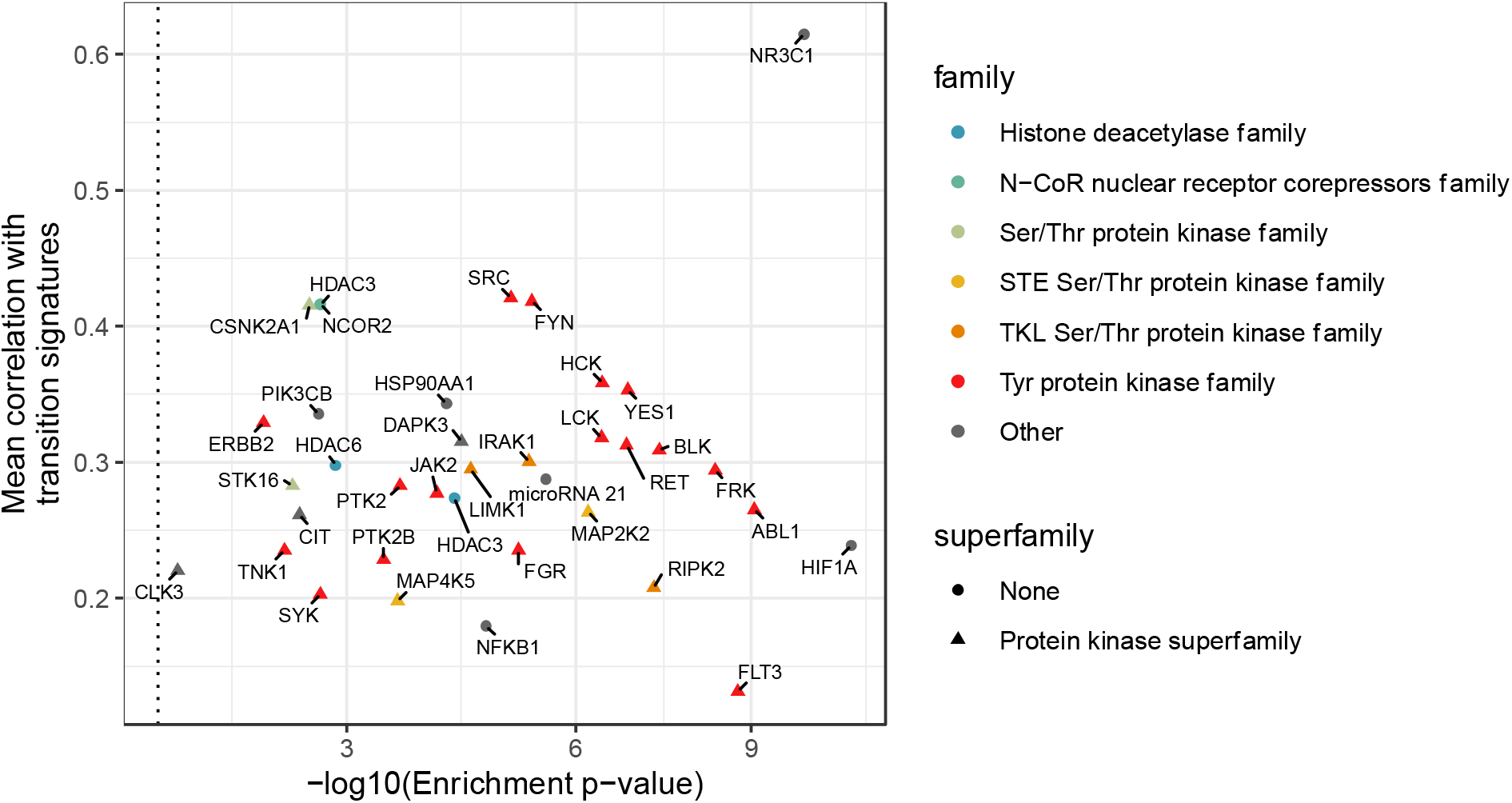
Correlation and enrichment of targets. For all targets with positive correlations between bioactivity and signature matching for both signatures and significance for at least one, the mean correlation for both signatures is shown, as well as the enrichment FDR which describes if high bioactivity (pCHEMBL > 5) was over-represented among the matched compounds, compared to all other compounds with perturbation signatures in the A549 cell line. The most significant enrichment is found for HIF1A and the highest correlation for NR3C1.

To gain a broader overview on potentially involved targets, a less conservative filtering was applied in which only significance for one of the two transition signatures was required. This identified 37 targets, shown in Figure 7, with HIF1A showing the strongest enrichment of bioactivity (pCHEMBL ≥ 5), while the strongest correlation was found for NR3C1. Multiple targets already implied in the previously identified transcriptional clusters were recovered (Figure 4), namely NR3C1, HSP90AA1, two histone deacetylases (HDAC3 and HDAC6) and the histone deacetylase 3 and nuclear receptor corepressor 2 protein complex (HDAC3/NcoR2). Furthermore, 24 protein kinases were identified including 17 tyrosine kinases as well as the casein kinase CSNK2A1, which was also inferred in upstream signalling based on the transition signatures (Figure 3). These were also found to be frequently co-identified, which can be explained by the known polypharmacology of kinase inhibitors ^385^ (Figure S 4). Additional targets included phosphatidylinositol-4,5-bisphosphate kinase catalytic subunit beta (PIK3CB) and miRNA 21 which was the only identified target which was not a protein or protein complex, and is a pleiotropic miRNA involved in pulmonary remodelling ^90^. Hence, this less stringent filtering is able to suggest additional targets, and recovers the primary targets linked to the previously transcriptional clusters.

To gain a more complete view on how the inferred targets may interact, we next derived physical interactions between the identified targets from Omnipath ^47^, as shown in Figure 8. Many interactions were found between the kinases with SRC followed by focal adhesion kinase (PTK2) showing the highest degree centrality, indicating that modulation of these may also linked to the modulation of many other proposed targets. However, here it should be noted that this underestimates the role of HDACs and HSP90, as these do not primarily act by interacting with or modulating individual proteins but by modulating a wide range of entities through indirect effects, namely, respectively, by epigenetically modulating their expression ^78^ and by stabilizing or degrading them ^83^. Although not all types of functions are described equally well, protein-protein interactions may provide useful insights into targets with central functions.

**Figure 8:**
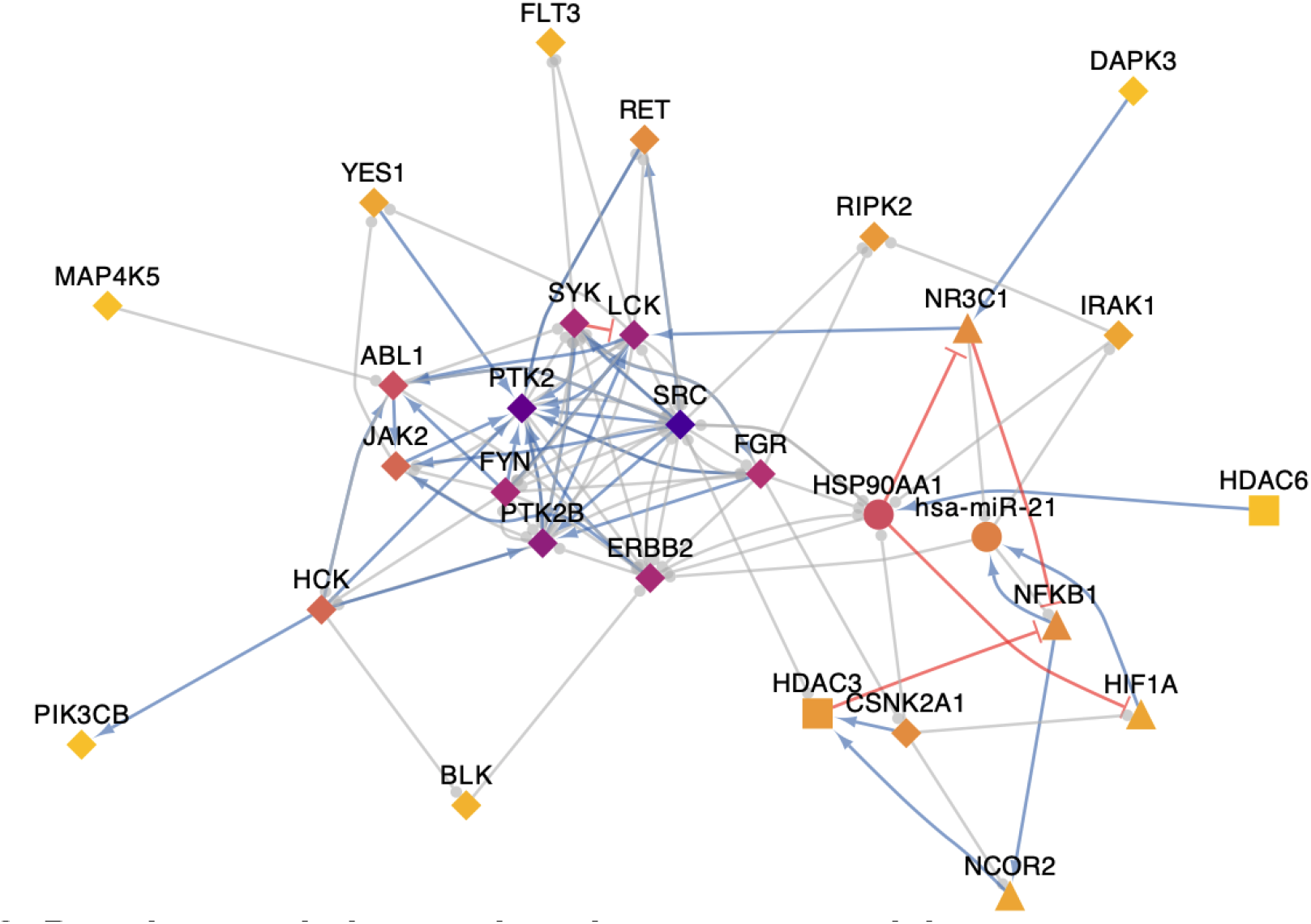
Protein-protein interactions between potential targets. Physical protein-protein interactions derived from Omnipath between potential targets are shown. Activatory (red) and inhibitory (blue) edges are indicated as colour, and edges without sign information are shown in grey. The highest degree centrality, indicated as node colour, is found for SRC followed by Focal adhesion kinase 1 (PTK2). Transcription factors (▲), epigenetic regulators (■), and kinases (⍰) are indicated through distinct node shapes given the specific interest in these targets in this study.

## Conclusion

Reinstating endogenous alveolar regeneration is an emerging therapeutic target in pulmonary fibrosis, due to its potential ability to not only reduce disease progression but to reverse it. Thanks to the advances in single-cell technologies, it has been possible to characterise the molecular identity of stem cell states involved in regeneration as well as their trajectories, and this has also led to the discovery of an intermediate progenitor cell state AT2 to AT1 cell differentiation in alveolar regeneration which is at the centre of this study. However, how to modulate these tightly regulated processes is not yet clear, also because key targets and mechanisms have yet to be discovered in the context of human lung disease.

In this study, we aim to computationally prioritize small molecules with the ability to promote the intermediate progenitor to AT1 cell transition. To this end, we use publicly available scRNA-Seq datasets, based on which the transition was initially discovered, to characterise the transcriptional changes associated with the transition and match this to compounds from the LINCS database which induce similar transcriptional changes and hence may promote the transition. Using this approach, we identify fostamatinib as most promising candidate which aligns with other studies which have prioritised it for acute lung injury ^75^ and showed that it improves the clinical outcome in COVID-19 ^76^. Overall, we identify multiple kinase inhibitors, glucocorticoid agonists, HDAC inhibitors and HSP90 inhibitors as matched, and also found that bioactivity for the respective targets of these compound classes was also found to correlate with signature matching providing complementary evidence that these targets are mechanistically linked to the cell transition (Figure 7).

As additional promising targets, NFKB1 and HIF1A are identified which are key TFs based on the transition signatures (Figure 3, Figure 5), and show a correlation between bioactivity and signature matching for both transition signatures (Figure 6). Furthermore, HIF1A also showed the most significant enrichment of activity (pCHEMBL ≥ 5) in matched compounds (Figure 6) and JUN was identified as most significantly differentially regulated TF (Figure 5). Indeed, termination of NKFB1-mediated inflammation ^86^, as well as hypoxia mediated by HIF1A ^87^, have previously been implicated in alveolar regeneration, and Choi *et al*. specifically demonstrated that the transient activation of NFκB and HIF1A is essential for the AT2 to AT1 transition ^16^.

Among the kinases, we find that SRC kinase family is particularly well represented, with three members being among the six targets which correlate to both transition signatures (Figure 6). Overall, SRC is also the most central node in the PPI network connecting potential targets, followed by PTK2 (Figure 8).

While the effect of the tested compounds on alveolar regeneration remains to be validated experimentally, this study hence demonstrates how scRNA-Seq data can be used for computational drug repurposing and how it is then possible to further prioritise potentially involved targets and downstream TFs. It should be highlighted, that similar cases where transitions are first understood transcriptionally can be expected to arise, thanks to single-cell transcriptomics. In these cases, signature matching might provide candidates for drug repurposing and, on a more general level, valuable starting points for drug discovery, also when additional information is not available yet.

### Appendix

**File S1: Cell annotations provided by Choi et al**.

The anno_final_v3 cluster annotation provided by Choi et al. was used to generate the celltype annotations used in Chapter 5. The file can be accessed via zenodo (DOI:10.5281/zenodo.7017239).

**Figure S 1:**
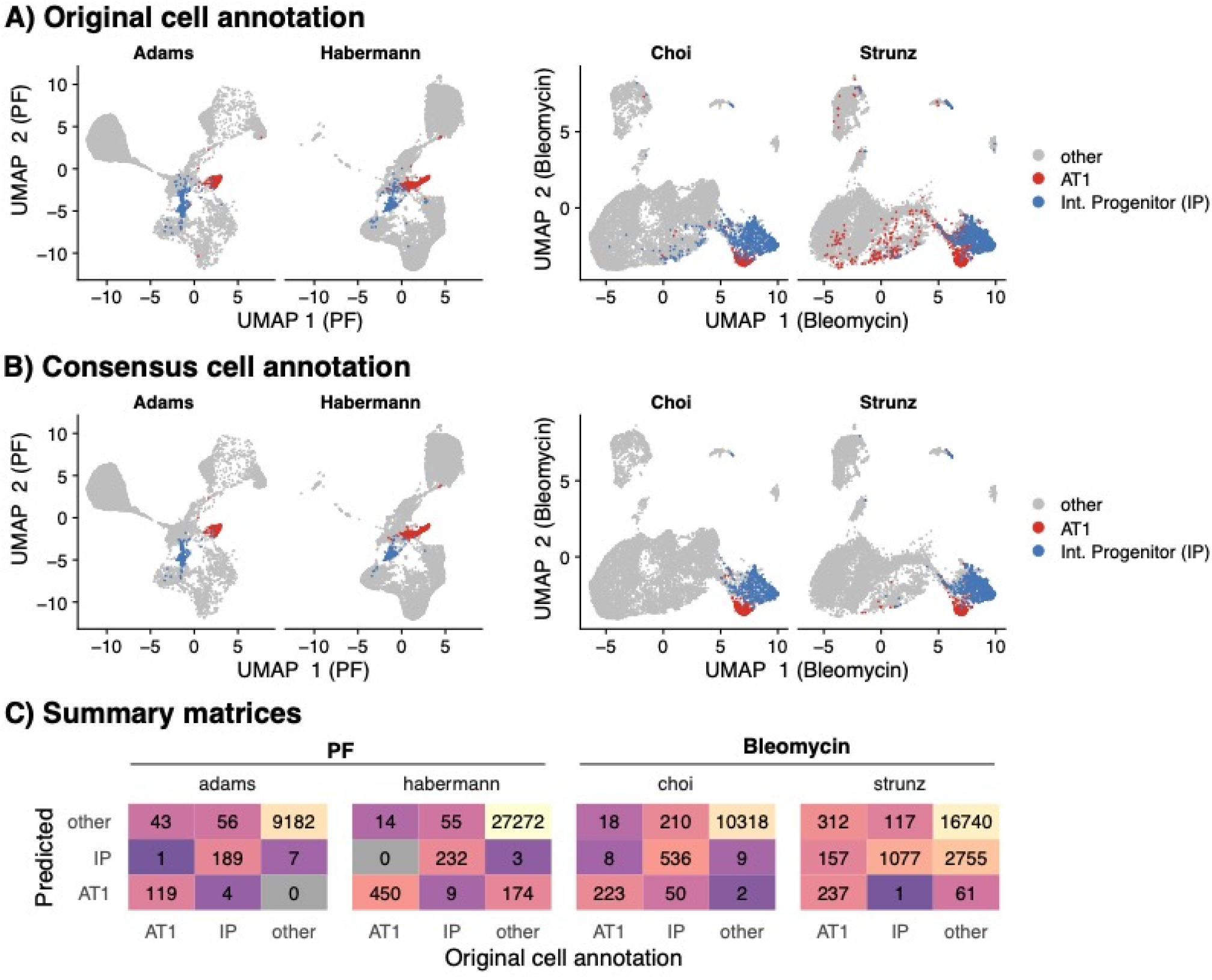
Comparison of original and consensus annotations for AT1 cells and intermediate progenitors (IP). A) Original cell annotations for each dataset plotted onto the UMAP after Harmony integration. PF datasets were first subsetted to epithelial non-ciliated cells to improve integration. B) Same UMAP projection with only consensus AT1 and IP cells indicated, which show matching original and predicted cell labels. C) Comparison of original and predicted labels.

**Figure S 2:**
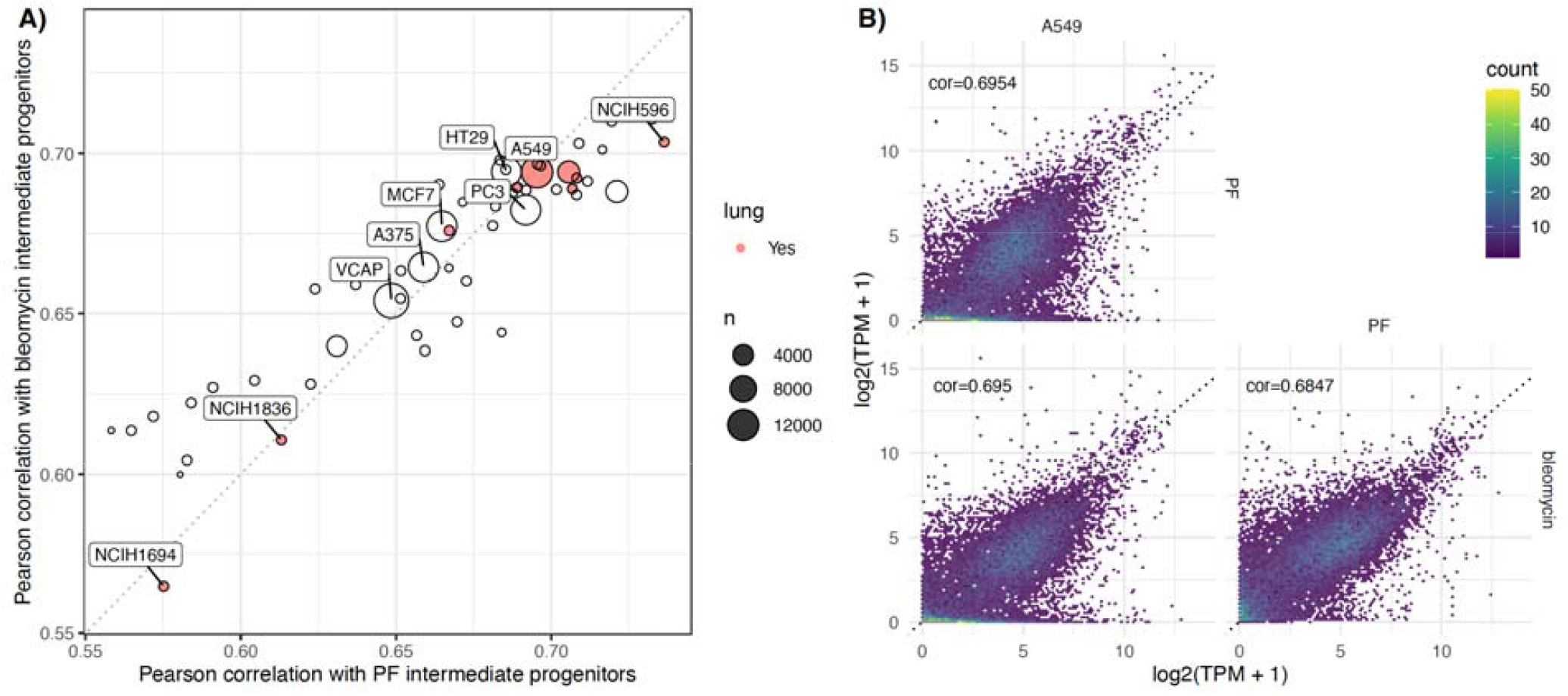
Comparison of baseline transcriptional profiles between LINCS cell lines and intermediate progenitor cells. The correlation between the baseline expressions of the intermediate progenitor population from IPF or bleomycin, respectively, and each cell line in LINCS is shown, and the number of perturbation signatures is shown as point size. Among the cell lines with at least 10,000 measured perturbations, the highest correlation is found for the A549 lung adenocarcinoma cell line. The highest correlation among cell lines of lung origin (red) is found for the NCIH596 adenosquamous carcinoma cell line, while the lowest ones are found for the only two small cell lung cancer (SCLC) cell lines NCIH1694 and NCIH1836. B) Comparison between the baseline expression of the A549 cell line and the intermediate progenitor populations reveals that the A549 cell line is as transcriptionally similar to the intermediate progenitor populations as they are to each other. All correlations were significant (p-value < 2.2e-16).

**Figure S 3:**
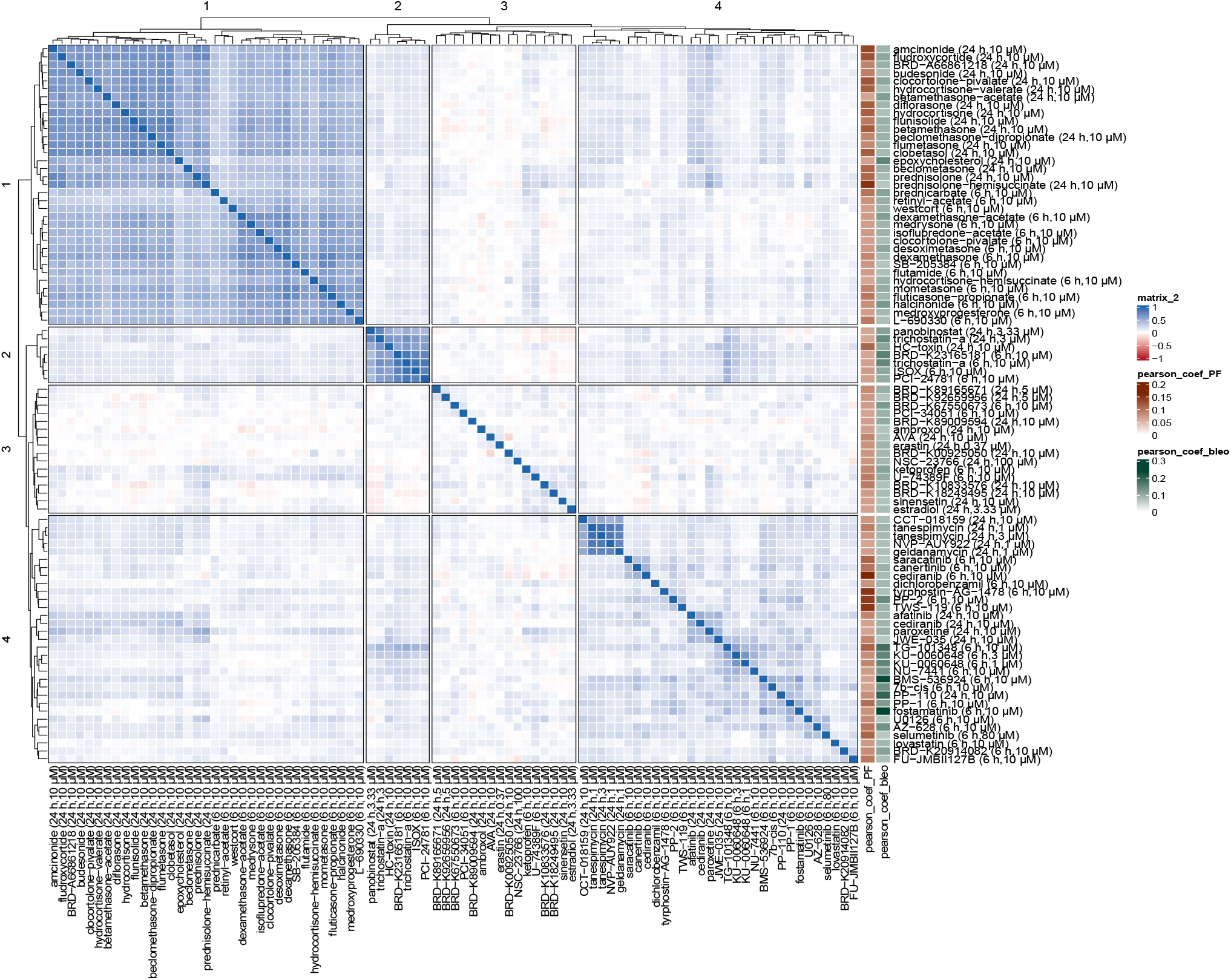
Pearson correlation between signatures of matched perturbations. The correlation between the consensus signatures for landmark genes is shown identifying multiple transcriptional compound clusters.

**Figure S 4:**
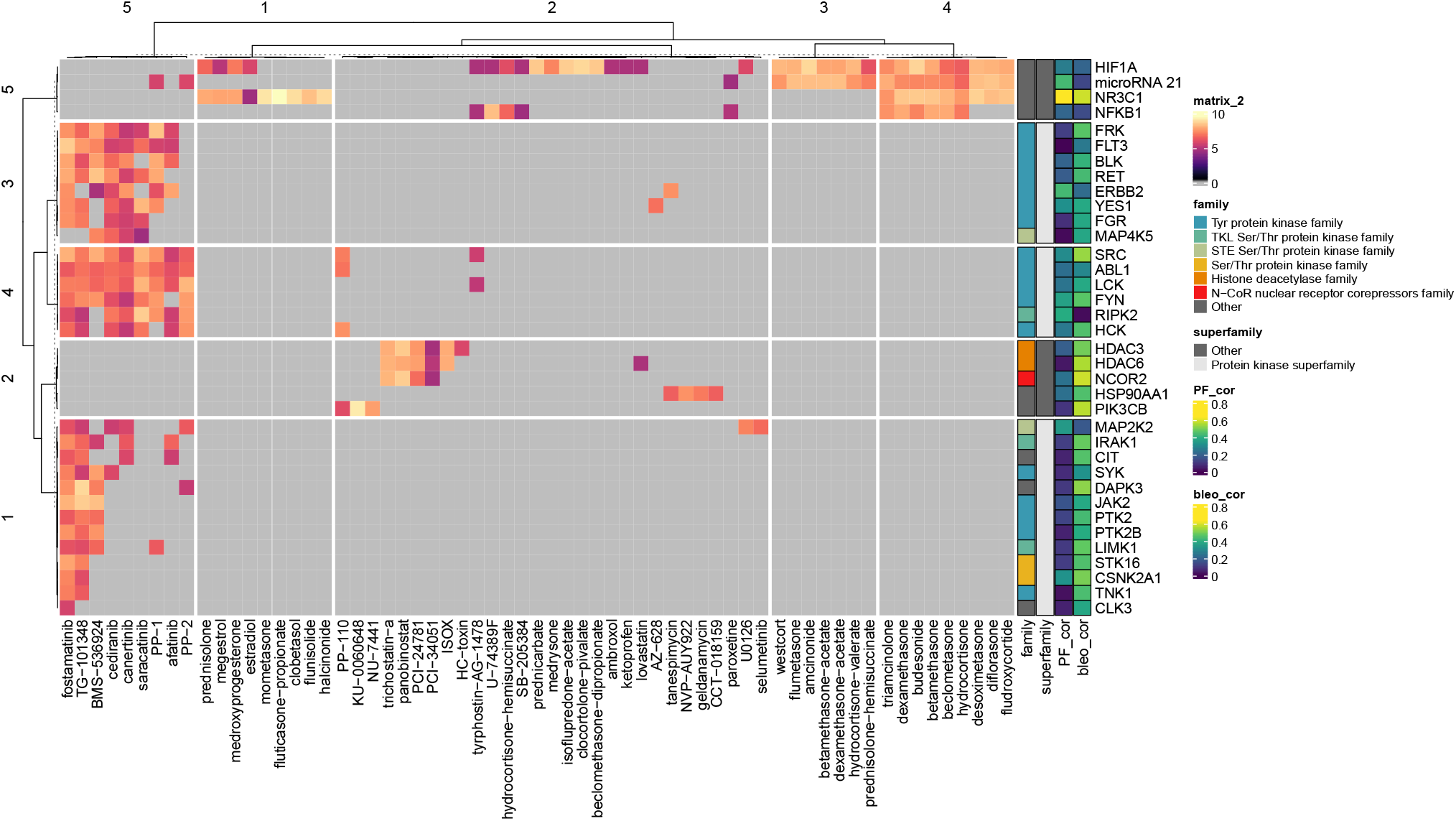
Bioactivity data for correlated targets and matched compounds. For all targets with positive correlations between bioactivity and signature matching for both signatures and significance for at least one (p-value < 0.05), the pCHEMBL values across matched compounds are shown and have been clustered using the Jaccard distance based on the presence or absence of a measured pCHEMBL value.

